# Diagnostic Value of Antibody Responses to *Mycobacterium avium* subsp. *paratuberculosis*-Derived Proteins PtpA and PtpB in Rheumatoid Arthritis

**DOI:** 10.64898/2026.01.08.698458

**Authors:** Jorge Hernández-Bello, Sergio Cerpa-Cruz, Gabriela A. Sánchez-Zuno, Ferdinando Nicoletti, Horacio Bach, José F. Muñoz-Valle

## Abstract

Evidence suggests that *Mycobacterium avium* subspecies *paratuberculosis* (MAP) may contribute to autoimmune diseases such as rheumatoid arthritis (RA), partly through effector proteins—particularly the tyrosine phosphatases PtpA and PtpB—that modulate macrophage signaling and promote bacterial persistence. This study evaluated whether serum antibodies against these proteins serve as biomarkers of RA. Humoral responses to PtpA and PtpB were quantified in Mexican RA patients (n = 100) and healthy controls (n = 100) using in-house ELISAs. Associations with disease activity (DAS28), ROC performance, and logistic regression models were assessed. Results showed that anti-PtpB antibody levels were significantly higher in patients with RA than in healthy controls (median OD 0.185 vs. 0.080; p < 0.0001) and had moderate discriminative capacity (AUC = 0.762). Anti-PtpB reactivity increased with higher disease activity and showed a significant positive association with DAS28 (p < 0.05). In addition, there was a functional disability measured by HAQ (p < 0.001), as well as moderate correlations with erythrocyte sedimentation rate and rheumatoid factor. A combined logistic regression model integrating both antibodies markedly improved diagnostic accuracy (AUC = 0.934), achieving high sensitivity (90%) and specificity (89%). These findings support a potential role of MAP in RA immunopathogenesis and indicate that combined quantification of anti-PtpA and anti-PtpB antibodies captures complementary and non-redundant immunological information. This combined serological approach may enhance RA diagnosis and provide clinically relevant insights into disease activity and severity.

## Introduction

RA is a multifactorial autoimmune disease in which genetic predisposition, dysregulated immune pathways, and microbial exposures interact to promote chronic synovial inflammation and joint destruction [1,2]. Growing interest has focused on microorganisms capable of persisting within host immune cells and generating antigenic stimuli that may shape autoantibody production or amplify inflammatory cascades. Among these, *Mycobacterium avium* subsp*. paratuberculosis (*MAP) has emerged as a plausible environmental trigger of autoimmunity due to its ability to survive within macrophages and modulate intracellular signaling via secreted virulence factors [3–5].

MAP effector proteins involved in host–pathogen interactions have attracted particular attention for their immunogenic properties and potential relevance in RA. A pivotal study from Italy demonstrated that the MAP-derived protein tyrosine phosphatases PtpA and PknG are recognized at significantly higher frequencies in the sera of RA patients than in those of healthy controls, supporting the notion that MAP exposure may leave a detectable humoral footprint in RA [6]. Building on this idea, our group recently reported that PtpA-specific antibodies are also elevated in Mexican patients with RA and may serve as an informative immunological marker in this population [7]. These observations collectively suggest that MAP phosphatases and kinases are antigenic targets that can elicit differential immune responses in RA.

Mechanistically, MAP-secreted proteins such as PtpA and PtpB may become relevant to RA pathogenesis because their intracellular effects converge on processes central to joint inflammation. PtpA inhibits phagolysosomal fusion by dephosphorylating the human VPS33B, a protein involved in phagolysosome fusion. This dephosphorylation allows bacteria to persist within macrophages [8], major producers of the pro-inflammatory cytokines TNF, IL-1β, and IL-6, which drive synovitis and structural damage [9]. Persistent MAP antigens could therefore act as chronic stimuli, maintaining macrophage activation, promoting continuous cytokine release, and enhancing Th1/Th17 polarization [10]. In genetically susceptible individuals, repeated exposure to these antigens may also increase autoantibody formation via molecular mimicry [11]. Furthermore, a higher antigenic load or stronger immune recognition of MAP phosphatases may reflect ongoing innate immune activation, potentially explaining an association between elevated antibody levels and greater clinical activity in RA.

Among MAP-secreted effectors, the tyrosine phosphatases PtpA and PtpB have attracted attention for their ability to disrupt phagosomal maturation and phosphoinositide metabolism, thereby interfering with vesicular trafficking and promoting intracellular survival—mechanisms that have been extensively characterized in related mycobacterial pathogens. Although evidence has begun to accumulate for PtpA-specific responses in RA, whether PtpB elicits a similar or complementary humoral signature remains unknown. Furthermore, whether combining immune responses to both phosphatases can improve diagnostic discrimination has never been explored.

In this study, we analyzed antibody reactivity to PtpA and PtpB in patients with RA and healthy controls and assessed the diagnostic utility of each marker individually and in combination. By integrating our previous findings [7] with new data on PtpB, we aimed to clarify the immunological relevance of MAP-secreted phosphatases in RA and determine whether their combined measurement enhances diagnostic accuracy.

## Materials and methods

### Subjects

Archived serum samples from patients and healthy controls were used in this study. These samples were used to determine the level of anti-PtpA in our previous study [7]. Briefly, RA patients (23 males, 77 females; median age 58) who fulfilled the 2010 ACR/EULAR Classification Criteria for RA were enrolled at the Rheumatology Unit of the Civil Hospital of Guadalajara, Fray Antonio Alcalde, Guadalajara, Jalisco, Mexico, between January 1, 2018, and December 31, 2021. Clinical and demographic data were collected, including disease duration, rheumatoid factor (RF) and anti-cyclic citrullinated peptide (anti-CCP) status, treatment with steroids and disease-modifying anti-rheumatic drugs (DMARDs), C-reactive protein (CRP) levels, erythrocyte sedimentation rate (ESR), Disease Activity Score-28 (DAS-28), and Health Assessment Questionnaire (HAQ) scores.

A group of 100 healthy controls (20 males, 80 females; median age 40 years) was recruited at the same hospital. Control participants verbally confirmed having no prior history of tuberculosis. Antibody reactivity to PtpA in this cohort has been previously described [7].

For the present study, the same archived sera were additionally evaluated for reactivity to PtpB, and combined analyses of PtpA and PtpB were performed.

### Ethics approval statement

This study was approved by the Ethics Committee of the University of Guadalajara (Approval No. 0122017) and conducted in accordance with the ethical principles outlined in the Declaration of Helsinki (64th World Medical Association General Assembly, Fortaleza, Brazil, 2013). Written informed consent was obtained from all participants prior to inclusion in the study.

### ELISA assays

#### Plate preparation

The recombinant PtpB protein was expressed in *Escherichia coli* harboring the *ptpB* gene in the ampicillin-resistant pET-22 vector. Purification was performed via Ni-NTA affinity chromatography, and the protein was stored at –20°C until use. The preparation of recombinant PtpA and the corresponding assay conditions have been previously described [7].

For ELISA, Maxisorp plates (ThermoFisher) were coated with 50 μg/mL of antigen in phosphate-buffered saline (PBS) and incubated overnight at 4°C. Plates were washed three times with PBS containing 0.05% Tween-20 (PBS-T) and blocked with 3% bovine serum albumin (BSA) in PBS at 4°C overnight. Plates were air-dried prior to use.

### Assay procedure

Sera from RA patients and controls were tested in triplicate. After incubation with sera, plates were washed with PBS-T and subsequently incubated with a peroxidase-conjugated anti-human IgG secondary antibody. Optical density (OD) was measured at 450 nm using an Epoch microplate reader (BioTek, USA). The baseline signal, defined as the secondary antibody alone, was subtracted from all readings. Positive controls were included in all assays. Cut-off values were determined by Receiver Operating Characteristic (ROC) analysis to ensure specificity above 90%, with sensitivity adjusted accordingly.

### Statistical analysis

Differences in antibody reactivity between RA patients and controls were assessed using the Mann–Whitney U test. Associations between clinical variables and antibody levels were examined by linear regression. ROC curves and areas under the curve (AUC) were generated using Python (version 3.14) with the scikit-learn library. A multivariable logistic regression model integrating anti-PtpA and anti-PtpB antibody levels was constructed to assess combined diagnostic performance. Pairwise comparisons between ROC curves were performed using DeLong’s test. Sensitivity, specificity, positive predictive value (PPV), and negative predictive value (NPV) were calculated at the optimal cutoff determined by the Youden index. Graphical representations of ROC curves and correlation plots were generated using GraphPad Prism version 8.0 (GraphPad Software, San Diego, CA). Statistical significance was defined as a two-sided p-value < 0.05.

## Results

Patients with RA were significantly older than controls (median 54 *vs*. 40 years, p < 0.0001), while the proportion of females was similar between groups (81% *vs*. 80%, p > 0.90). Among RA patients, disease activity and disability indices showed median DAS-28 and HAQ scores of 3.2 and 0.75, respectively. Inflammatory markers were elevated, with C-reactive protein (CRP) at 6 mg/dL and ESR at 20 mm/h. Most patients were receiving DMARDs (73%), and more than one-third reported NSAID use (36%), whereas only 1% were on corticosteroid therapy (Table 1).

**Table 1.** Clinical and demographic features of RA patients and control subjects (CS).

| Variable | RA (n = 100) | Controls (n = 100) | P-value |
| --- | --- | --- | --- |
| Age, years (median, IQR) | 54 (41–61) | 40 (31–53) | <0.0001 |
| Female sex, n (%) | 81 (81%) | 80 (80%) | >0.90 |
| HAQ score (0–3) | 0.75 (0.25–1.25) | — | — |
| DAS-28 score | 3.2 (2.6–5.1) | — | — |
| C-reactive protein (CRP), mg/dL | 6 (2–12) | — | — |
| Erythrocyte sedimentation rate (ESR), mm/h | 20 (12–44) | — | — |
| White blood cells (WBC), $\times 10^9/L$ | 7.0 (5.8–8.5) | — | — |
| Red blood cells (RBC), $\times 10^{12}/L$ | 4.5 (4.1–4.9) | — | — |
| Hemoglobin (Hb), g/dL | 13.4 (12.1–14.8) | — | — |
| Mean corpuscular volume (MCV), fL | 31.0 (29.0–33.0) | — | — |
| Platelets (PLT), $\times 10^9/L$ | 260 (210–320) | — | — |
| Rheumatoid factor (RF), IU/mL | 45 (20–118) | — | — |
| Body weight, kg | 65 (56–74) | — | — |
| Height, cm | 158 (152–165) | — | — |
| Body mass index (BMI), kg/m<sup>2</sup> | 27.5 (24.0–31.5) | 26.3 (23.6–32) | >0.90 |
| Disease evolution time, years | 8 (2–20) | – | – |
| Current steroid therapy, n (%) | 1 (1%) | – | – |
| DMARDs, n (%) | 73 (73%) | – | – |
| NSAIDs, n (%) | 36 (36%) | – | – |

Data are expressed as median (interquartile range, IQR) for continuous variables and as absolute numbers with percentages for categorical variables. Age was compared using the Mann–Whitney U test, and sex distribution was analyzed using the chi-square test. A $p < 0.05$ was considered statistically significant. Abbreviations: RA, rheumatoid arthritis; CS, control subjects; HAQ, Health Assessment Questionnaire; DAS-28, Disease Activity Score 28; DMARDs, disease-modifying antirheumatic drugs; NSAIDs, nonsteroidal anti-inflammatory drugs.

As shown in Fig 1A, antibody levels against PtpB were higher in the RA group than in controls (p < 0.0001, Mann–Whitney U test). The median optical density (OD) in RA patients was 0.1847 [25^th^–75^th^ percentile: 0.1120–0.2483], whereas the control group exhibited a median OD of 0.0801 [25^th^–75^th^ percentile: 0.03275–0.1434]. ROC analysis demonstrated that anti-PtpB antibodies effectively discriminated RA patients from healthy controls, with an AUC of 0.762 (p < 0.0001; Fig 1B).

**Fig 1.**
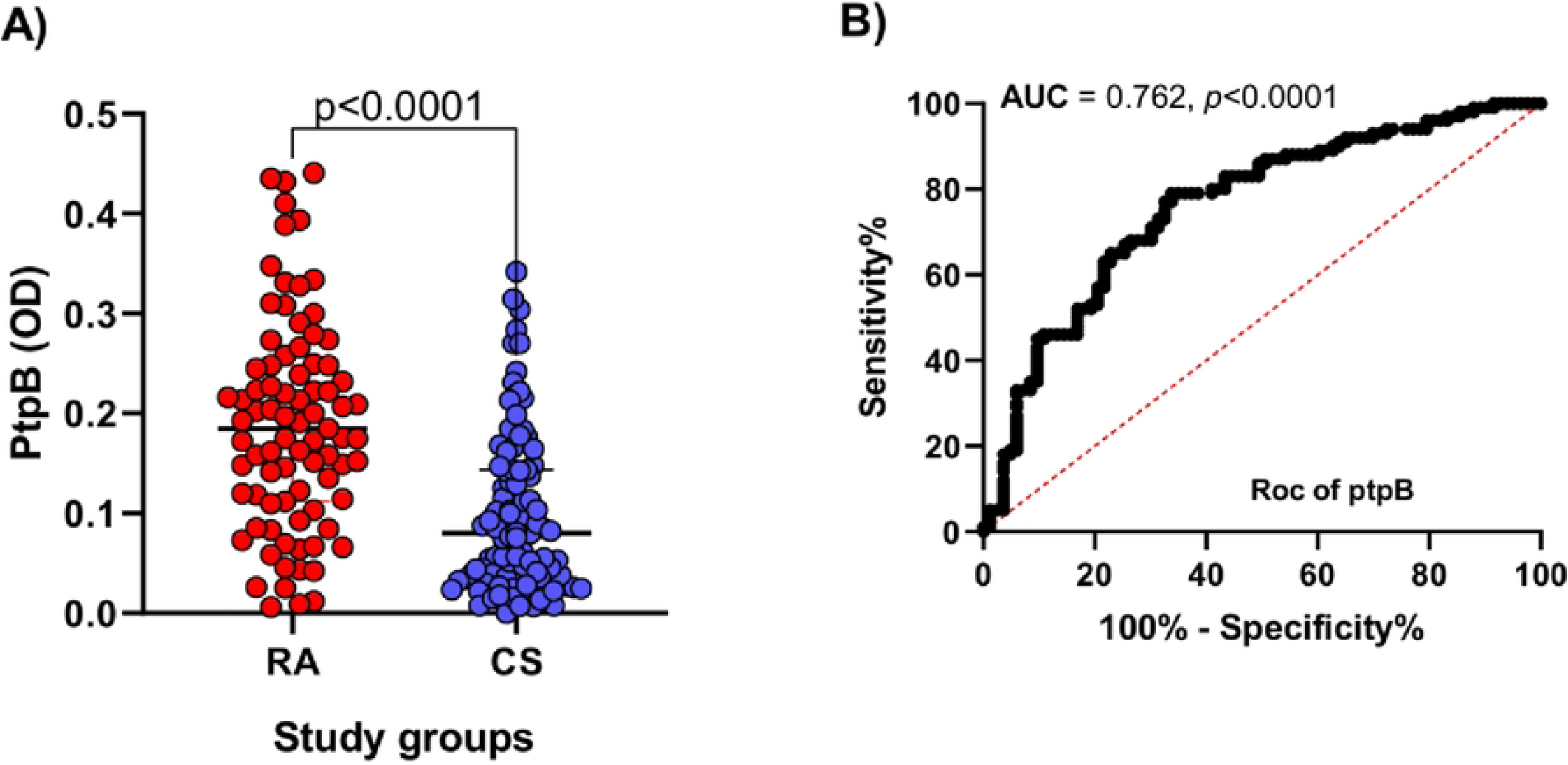
Serum antibody reactivity to PtpB in RA subjects and CS. (A) Comparison of anti-PtpB antibody levels between groups. Bars represent the median and interquartile range; dashed lines indicate antibody positivity thresholds, and p-values are shown above. (B) Receiver operating characteristic (ROC) curve evaluating the discriminative capacity of anti-PtpB antibodies.

To assess associations between anti-PtpB levels and disease activity, RA patients were stratified into four groups according to their DAS28 scores: remission, low disease activity, moderate disease activity, and high disease activity. As shown in Fig 2, patients in remission had the lowest antibody levels (median = 0.1187; IQR = 0.0447–0.1887). Those with low disease activity showed slightly higher values (median = 0.1752; IQR = 0.1053–0.2296), with no significant difference relative to remission (p = 0.3209). The moderate activity group displayed intermediate values (median = 0.1948; IQR = 0.1574–0.2527), which overlapped with those of the low activity group (p = 0.9999). Conversely, patients with high disease activity exhibited the highest antibody levels (median = 0.2357; IQR = 0.1682–0.3324) and differed significantly from the remission group (p = 0.0001).

**Fig 2.**
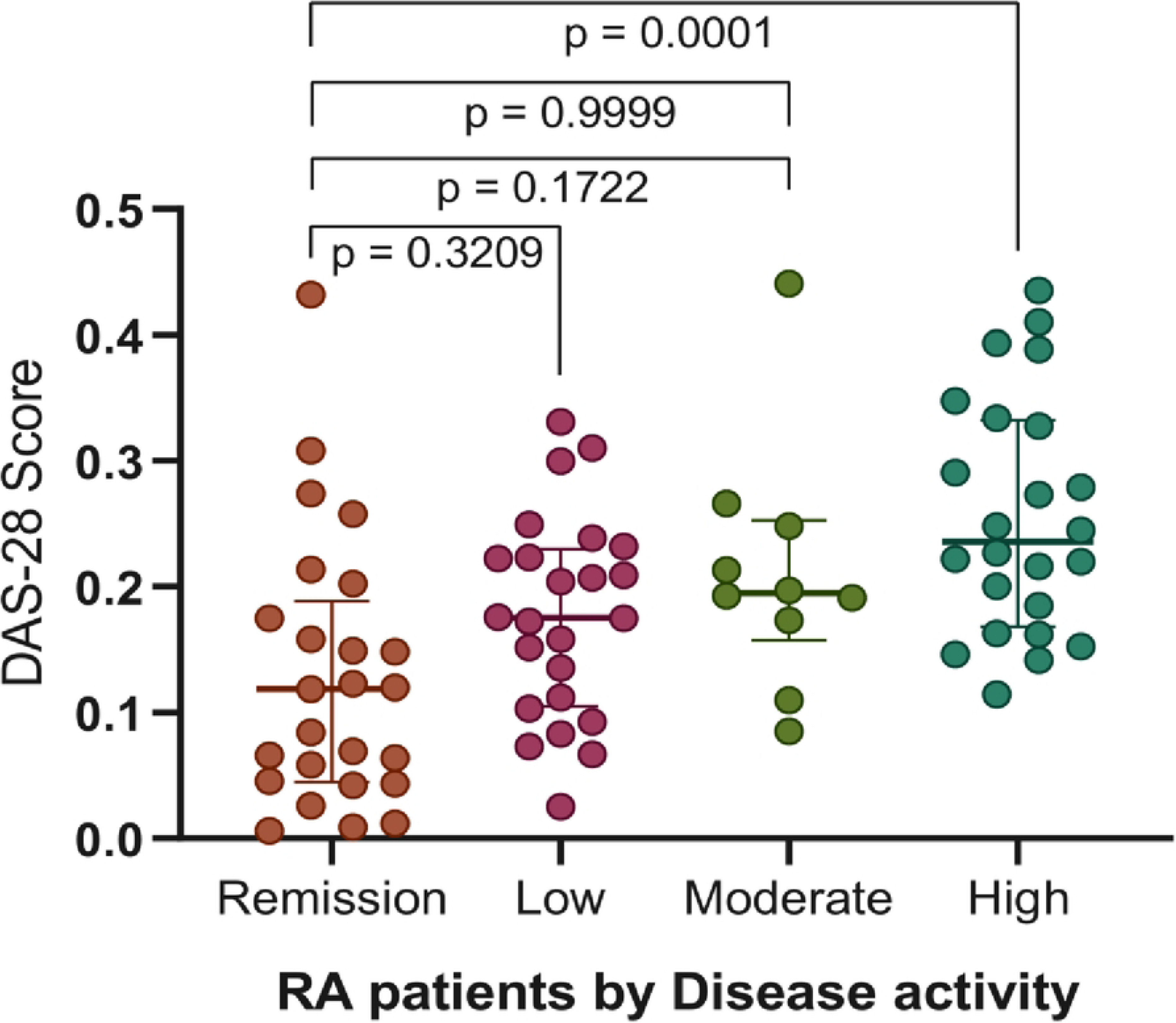
Association between serum anti-PtpB antibody levels and DAS-28 categories. Bars represent median ± interquartile range for each disease activity group. Statistical analysis was performed using the Kruskal–Wallis test followed by Dunn’s post hoc correction. P-values are shown above the distributions.

Fig 3 displays the correlation heatmap between anti-PtpB antibody levels and clinical and laboratory variables in patients with RA. Anti-PtpB antibody levels showed statistically significant positive correlations with disease activity and functional impairment, including DAS28 (ρ = 0.45, p < 0.001) and HAQ (p = 0.40, p < 0.001). In addition, moderate positive associations were observed with ESR (p = 0.37, p = 0.003) and rheumatoid factor (RF; p = 0.49, p < 0.001). No significant differences in anti-PtpB levels were observed by sex or treatment status, including use of non-steroidal anti-inflammatory drugs (NSAIDs), corticosteroids, sulfasalazine, chloroquine, or methotrexate. No other clinical, hematological, or demographic variables were significantly associated with anti-PtpB levels.

**Fig 3.**
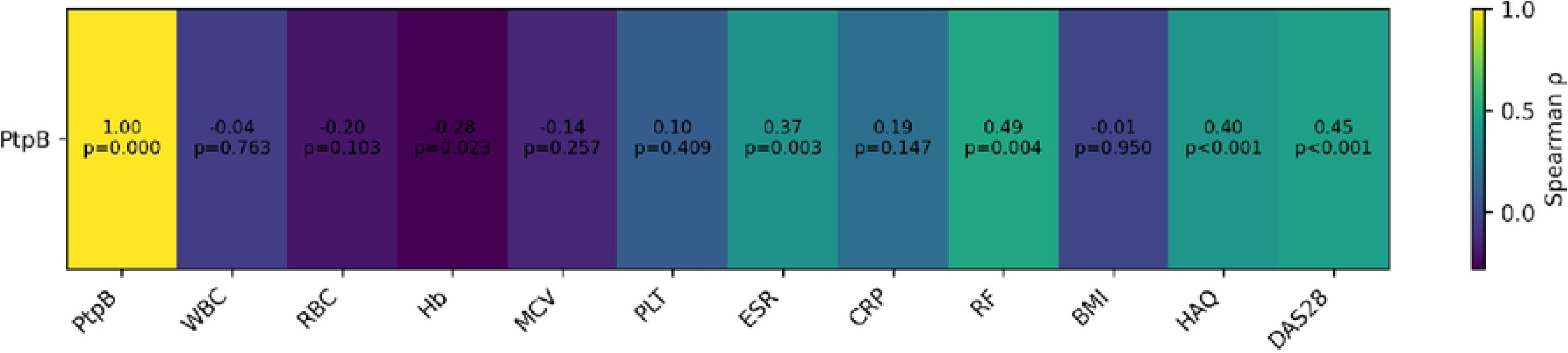
Correlation heatmap of anti-PtpB antibody levels with clinical and laboratory parameters in patients with RA. The heatmap displays Spearman correlation coefficients (ρ) between anti-PtpB antibody levels and clinical and laboratory variables, including hematological parameters, inflammatory markers, functional disability, and disease activity indices. Each cell shows the corresponding correlation coefficient and p-value. Warm colors indicate positive correlations, whereas cool colors indicate negative correlations. Abbreviations: WBC, white blood cells; RBC, red blood cells; Hb, hemoglobin; MCV, mean corpuscular volume; PLT, platelets; ESR, erythrocyte sedimentation rate; CRP, C-reactive protein; RF, rheumatoid factor; BMI, body mass index; HAQ, Health Assessment Questionnaire; DAS28, Disease Activity Score 28.

The relationship between humoral immune responses to the MAP-derived proteins PtpA and PtpB was assessed by comparing antibody levels in RA patients and controls. As shown in Fig 4A and 4B, no significant correlation was found between anti-PtpA and anti-PtpB levels in either group.

**Fig 4.**
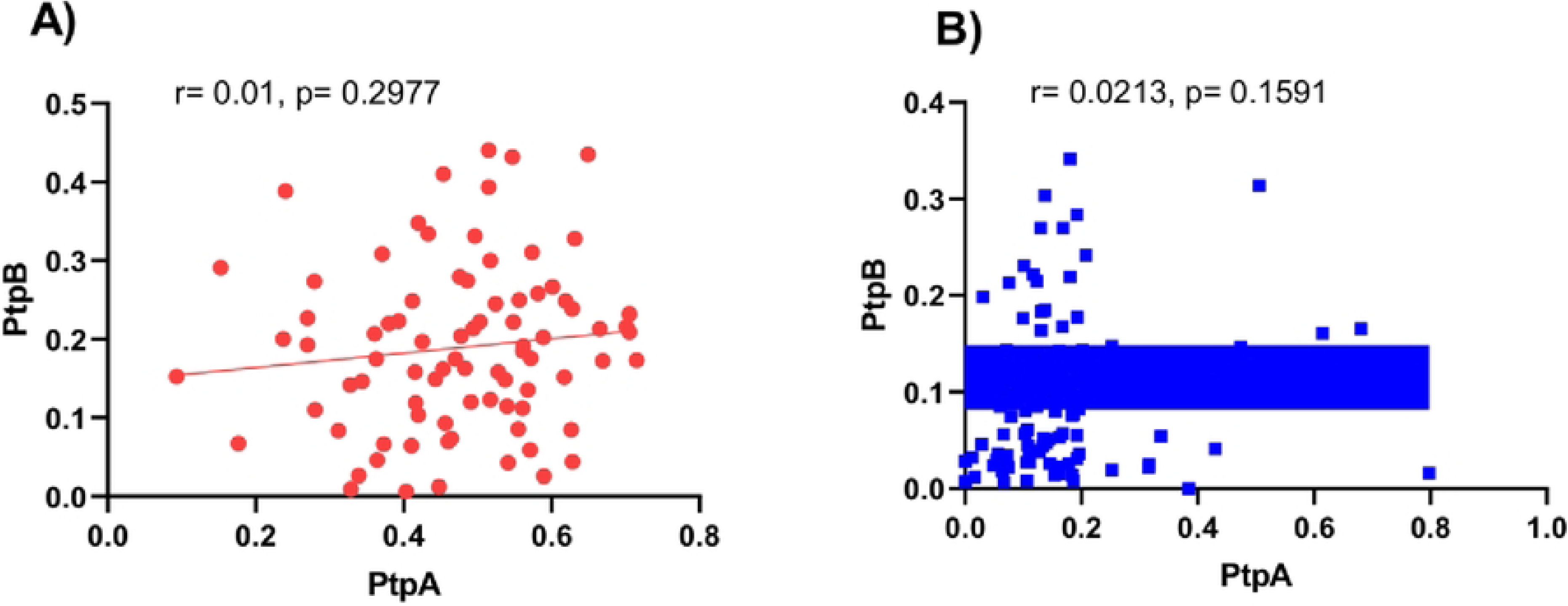
**Correlation between anti-PtPA and anti-PtpB antibody levels in RA patients and control subjects. (**A) RA patients. (B) Control subjects. Spearman’s rank correlation analysis was used to assess associations between markers. Blue and red lines denote the best-fit linear regressions for each group.

To evaluate the diagnostic utility of combining anti-PtpA and anti-PtpB antibodies, a multivariable logistic regression model was constructed and its performance assessed. As shown in Table 2, the combined model demonstrated excellent discriminative capacity, with an AUC of 0.934. At the optimal cut-off value of 0.3869, determined using the Youden index, the model achieved a sensitivity of 96% and a specificity of 87%. This threshold also yielded high predictive values, with a positive predictive value (PPV) of 86% and a negative predictive value (NPV) of 97%.

**Table 2.** Diagnostic performance of the logistic regression model combining anti-PtpA and anti-PtpB antibody levels.

| Metric | Value |
| --- | --- |
| Area under the curve (AUC) | 0.934 |
| Optimal cut-off point (Youden index) | 0.3869 |
| Sensitivity | 0.9638 |
| Specificity | 0.87 |
| Positive predictive value (PPV) | 0.8602 |
| Negative predictive value (NPV) | 0.9666 |

ROC curves for anti-PtpA, anti-PtpB, and their combination are presented in Fig 5. The combined model outperformed both individual markers (PtpA AUC = 0.925; PtpB AUC = 0.762). Pairwise comparisons of ROC curves using DeLong’s test revealed no statistically significant differences between the combined model and anti-PtpA alone (ΔAUC = 0.009, p = 0.975), nor between the combined model and anti-PtpB alone (ΔAUC = 0.172, p = 0.495). Similarly, the difference between anti-PtpA and anti-PtpB was not statistically significant (ΔAUC = 0.163, p = 0.485).

**Fig 5.**
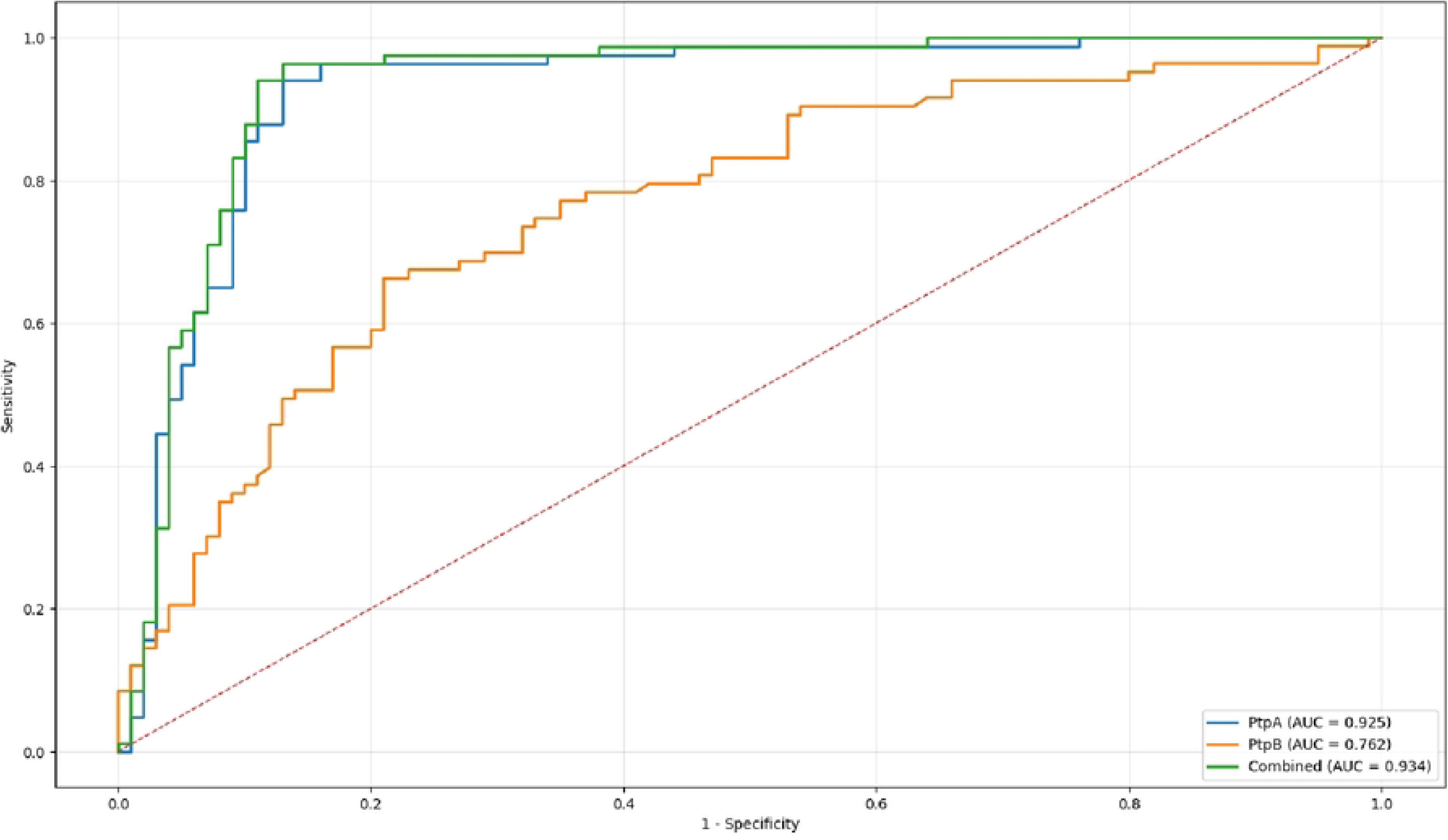
ROC curve comparison of models using anti-PtpA, anti-PtpB, and their combination. The green solid line represents the combined PtpA+PtpB model, which achieved the highest diagnostic accuracy. The blue dashed line represents the model using only PtpA, while the orange dashed line represents the model using only PtpB. The black diagonal line indicates random classification (AUC = 0.5).

## Discussion

Emerging evidence suggests that bacterial exposures may contribute to the etiology of RA by disrupting immune tolerance and promoting chronic inflammation in genetically susceptible individuals [18]. Several microorganisms, particularly those capable of persisting within macrophages or modifying host proteins, have been implicated as potential triggers of autoimmunity [19]. Intracellular bacteria such as MAP can interfere with phagosome maturation and sustain pro-inflammatory cytokine production [20,21]. Meanwhile, mucosal pathogens, including *Porphyromonas gingivalis* and *Aggregatibacter actinomycetemcomitans*, have been associated with aberrant protein citrullination and the induction of anti-CCP antibodies in RA [22], supporting the concept that microbial factors may converge on shared immunopathogenic pathways.

In this study, we evaluated humoral immune responses to the MAP-derived tyrosine phosphatases PtpA and PtpB in patients with RA and healthy controls. Four principal findings emerged: (i) anti-PtpB antibodies were significantly elevated in RA; (ii) anti-PtpB titers increased with higher disease activity; (iii) anti-PtpA and anti-PtpB responses were immunologically independent; and (iv) a combined logistic model incorporating both antibodies markedly improved diagnostic accuracy. Collectively, these observations strengthen the hypothesis that MAP antigens may contribute to the immunological landscape of RA and may serve as complementary biomarkers in this population.

Our findings extend existing evidence suggesting MAP exposure in RA. Previous studies have shown that MAP-secreted proteins, such as PtpA and PknG, are more frequently recognized by RA sera than by healthy control sera [12]. We also previously reported increased anti-PtpA responses in Mexican RA patients [7]. Molecular investigations in the USA, Europe, and the Middle East have detected MAP DNA or MAP-reactive antibodies in RA populations, reinforcing the plausibility of MAP as an environmental contributor to autoimmunity [6,23–25].

The present study adds the observation that PtpB, an established virulence phosphatase in mycobacteria, also elicits increased antibody responses in RA. Given that PtpB participates in immune evasion and intracellular persistence [16,26], heightened seroreactivity in RA patients is biologically compatible with chronic antigen exposure. Importantly, anti-PtpB antibody levels were positively associated with established measures of disease activity and functional impairment, including DAS28 and HAQ, providing a clinically meaningful link between MAP-related immune responses and disease expression. The association between anti-PtpB antibody levels and clinical disease activity provides a biologically plausible link between MAP-related immune responses and established pathogenic mechanisms in RA. Macrophages play a central role in RA synovitis, acting as key effector cells that produce pro-inflammatory cytokines such as TNF-α, IL-1β, and IL-6, which drive both joint inflammation and structural damage [27]. MAP phosphatases directly modulate macrophage biology: PtpA inhibits recruitment of the vacuolar H⁺-ATPase to the phagosome, blocking acidification [28], whereas PtpB in other mycobacteria modulates intracellular kinase signaling and promotes bacterial survival [29,30]. In this context, elevated anti-PtpB titers in RA may reflect sustained activation of innate immune pathways, consistent with their correlations with DAS28 and HAQ.

Anti-PtpB levels were also moderately associated with ESR and RF, but not with CRP or BMI. This pattern suggests that anti-PtpB immunity does not simply mirror acute-phase inflammation or metabolic status. Rather, it supports the existence of a more specific immunological axis, potentially centered on macrophage activation and humoral autoimmunity, in which MAP-related antigens contribute to disease severity without acting as nonspecific inflammatory markers. The lack of association with CRP may further indicate that anti-PtpB antibodies capture chronic or cumulative immune activation rather than transient inflammatory fluctuations.

The absence of significant differences in anti-PtpB levels according to sex or exposure to conventional antirheumatic therapies adds an important dimension to this interpretation. These findings suggest that anti-PtpB responses are relatively stable and not readily modulated by therapies that primarily target downstream inflammatory cascades, raising the possibility that anti-PtpB antibodies reflect upstream or parallel pathogenic processes that are not fully addressed by current therapeutic strategies.

One of the most distinctive findings was the independence between anti-PtpA and anti-PtpB titers in both RA patients and controls. This observation is consistent with and provides mechanistic support for our previous findings that anti-PtpA antibody levels were not associated with DAS28, autoantibody status, or inflammatory markers in RA [7]. Such independence likely reflects the functional divergence between the two phosphatases. PtpA disrupts the VPS33B–V-ATPase axis, primarily affecting phagosomal maturation and intracellular trafficking [31]. In contrast, PtpB operates through distinct lipid-mediated and kinase-dependent signaling pathways that influence host immune activation [32,33]. Their non-overlapping virulence mechanisms provide a plausible biological basis for differential immune recognition, suggesting engagement of distinct antigen-processing routes and B-cell activation pathways rather than a shared humoral response. In this framework, anti-PtpA responses may reflect exposure-related or host–pathogen interactions, whereas anti-PtpB immunity appears more closely linked to clinically relevant inflammatory and disease-activity pathways.

The strong diagnostic performance of the combined anti-PtpA/anti-PtpB logistic regression model further supports this complementary behavior. The model achieved excellent discriminative accuracy (AUC = 0.934), a level conventionally interpreted as indicative of high diagnostic utility [34]. Although pairwise ROC comparisons did not demonstrate statistically significant superiority over individual antibodies, the combined model consistently yielded numerically higher performance metrics. At the Youden-optimized cut-off, the model achieved very high sensitivity (96%) and negative predictive value (97%), while maintaining favorable specificity (87%) and positive predictive value (86%). This diagnostic profile is comparable to that reported for multiepitope serological tools used in early RA [35,36], underscoring the potential of MAP-derived antigens as clinically meaningful complementary biomarkers rather than standalone diagnostic replacements.

Accordingly, PtpA/PtpB serology is not intended to replace established RA biomarkers such as RF or anti-citrullinated protein antibodies, which remain central to the ACR/EULAR classification criteria. Instead, these MAP-derived immune markers may provide orthogonal information related to disease biology and immune activation. Significantly, future studies should extend the evaluation of these markers to other autoimmune and inflammatory diseases to determine their disease specificity.

MAP has been implicated in autoimmune diseases through mechanisms of molecular mimicry, including shared epitopes between MAP Hsp65 and human GAD65 [37]. Although this study did not evaluate epitope overlap, cross-reactive immunity may contribute to the amplification of adaptive responses, including the formation of RA-associated autoantibodies [38].

While this study was not designed to investigate transmission routes, previous work has demonstrated that MAP can be acquired through consumption of unpasteurized dairy products or contact with infected livestock, both of which have been associated with increased MAP positivity in humans. Given that MAP is shed in milk, feces, and aerosols from infected ruminants, these findings underscore the importance of zoonotic and foodborne exposure pathways in interpreting MAP-derived immune responses [39,40]. Such reservoirs may be particularly relevant in regions where traditional dairy practices persist or where rural populations have greater exposure to livestock. Future studies should therefore incorporate detailed exposure histories. In addition, the influence of long-term immunosuppressive therapies should be carefully considered, as disease-modifying antirheumatic drugs may alter host immune surveillance.

Some limitations should be considered when interpreting these findings. The cross-sectional design does not allow causal relationships to be established, and direct detection of MAP in tissues was beyond the scope of the present study. Although age differences were observed between groups, age did not correlate with antibody titers, suggesting that cumulative environmental exposure is unlikely to account for the observed MAP-reactive immune responses. Together, these data provide a strong rationale for future longitudinal and mechanistic studies to define further the pathogenic relevance of MAP-derived immune signatures in RA and their potential utility for patient stratification.

## Conclusions

This study demonstrates that anti-PtpB antibodies are elevated in RA, are associated with disease activity and functional impairment, and provide clinically relevant information that complements anti-PtpA responses. While anti-PtpA remains the strongest individual discriminator between RA patients and healthy controls, anti-PtpB antibodies appear to capture a distinct immunological dimension linked to inflammatory burden and disease severity. Accordingly, the combined assessment of both antibodies achieves excellent overall diagnostic performance, reflecting their complementary and non-redundant biological roles. Importantly, anti-PtpB antibody levels were independent of sex and conventional antirheumatic treatments, supporting the notion that MAP-related immune responses are not merely secondary to therapy or demographic factors. Together, these findings reinforce the hypothesis that exposure to MAP may represent a relevant environmental component in RA immunopathogenesis, with different MAP-derived antigens contributing heterogeneously to disease expression.

Further studies incorporating longitudinal sampling, tissue-level MAP detection, treatment stratification, and host genetic factors are warranted to clarify the mechanistic and clinical significance of MAP-related immunity in RA. Such efforts will also be essential for determining the specificity and broader relevance of combined anti-PtpA/anti-PtpB profiling in other autoimmune and rheumatic diseases in which MAP has been proposed as a potential environmental trigger.

## Acknowledgments

The authors would like to thank the patients who participated in this study.

## Author contributions

**JHB**: Formal analysis, Investigation, Methodology, Validation, Visualization, Writing – original draft, Writing – review and editing. **HB**: Conceptualization, Data curation, Formal analysis, Investigation, Methodology, Validation, Visualization, Writing – original draft, Writing – review and editing. **SCC**: Formal analysis, Investigation, Validation, Writing – original draft, Writing – review and editing. **GSZ**: Formal analysis, Investigation, Methodology, Validation, Visualization, Writing – review and editing. **FN**: Formal analysis, Investigation, Methodology, Validation, Visualization, Writing – review & editing. **FMV**: Conceptualization, Data curation, Formal analysis, Investigation, Methodology, Validation, Visualization, Writing – original draft, Writing – review and editing.

## Data availability

Data will be made available upon reasonable request.

## Supporting information

S1 Data. (XLSX)

